# Application of ATAC-Seq for genome-wide analysis of the chromatin state at single myofiber resolution

**DOI:** 10.1101/2021.06.14.448380

**Authors:** Korin Sahinyan, Darren M. Blackburn, Marie-Michelle Simon, Felicia Lazure, Tony Kwan, Guillaume Bourque, Vahab D. Soleimani

## Abstract

Skeletal myofibers are the main components of skeletal muscle which is the largest tissue in the body. Myofibers are highly adaptive in nature and they can vary in different biological and disease conditions. Therefore, transcriptional and epigenetic studies on myofibers are crucial to discover how chromatin alterations occur in the skeletal muscle under different conditions. However, due to the heterogenous nature of skeletal muscle, studying myofibers in isolation proves to be a challenging task. Single cell sequencing has permitted for the study of the epigenome of isolated myonuclei. While this provides sequencing with high dimensionality, the sequencing depth is lacking, which makes comparisons between different biological conditions difficult. Here we report the first implementation of single myofiber ATAC-Seq, which permits for the sequencing of an individual myofiber at a depth sufficient for peak calling and for comparative analysis of chromatin accessibility under various physiological, physical and disease conditions. Application of this technique revealed significant differences in chromatin accessibility between resting and regenerating myofibers. This technique can lead to wide application in identifying chromatin regulatory elements and epigenetic mechanisms in muscle fibers during development and in muscle-wasting diseases.

## INTRODUCTION

Skeletal muscle evolved for contraction and production of force. The main component of skeletal muscle are myofibers which are formed from the fusion of myogenic precursor cells (1) and myofibers are large postmitotic syncytia that are composed of repeating contractile units, called sarcomeres (2). Myofibers exhibit wide variations in their metabolic activity and contractile properties (3). In addition, they have a highly adaptive nature where their size, and the overall skeletal muscle mass, are regulated by complex processes involving rates of protein turnover (4, 5) transcriptional (6) and posttranscriptional (7) control of gene expression. Due to their adaptive nature, myofibers can change in response to exercise (3,8,9), aging (10) and diseases, such as sarcopenia (11, 12) and cachexia (13). Therefore, the study of the myofiber transcriptome/epigenome can provide key insights into how skeletal muscle adapts and changes under different conditions, and it can potentially lead to discovery of novel therapeutic venues for muscle related diseases.

Myofibers also act as a key signaling component of muscle stem cells (MuSCs) (14), which are in turn required for the regeneration of nascent muscle fiber after injury. Skeletal muscle is a very heterogenous tissue composed not only of myofibers and their associated MuSCs, but also numerous different non-myogenic cell types (15). Previous studies using whole muscle sequencing captures not only the myofibers but also the other resident cell types in the muscle, making it challenging to attribute changes in transcriptome and epigenome specifically to myofibers as they could be due to changes in other cell types. Recent advances in Next Generation Sequencing (NGS) now allows for high dimensional analysis at a single cell level. Recent studies using these technologies to study muscle tissue, such as single nucleus RNA-Seq and ATAC-Seq have analyzed the transcriptome and epigenome of the myonuclei within the muscle fiber (16–18). However, they present certain limitations where they sequence all myonuclei present in the muscle and cannot distinguish between different myofibers as well as having low sequencing depth with limited capacity of analysis.

Chromatin states play key roles in transcriptional regulation and determination of cellular identity (19). Although the accessible regions make up only 3% of the total genome, it represents over 90% of known transcription factor binding sites (20). Differential chromatin accessibility is a determinant of transcriptional gene regulation of different cell types and changes in chromatin accessibility have been shown in different biological and disease conditions such as during development (21, 22), cancers (23, 24) and neurological disorders (25, 26) among many others. Thus, in recent years, the study of epigenetics and chromatin accessibility has become a promising field for the development of novel therapeutics. Today, ATAC-Seq is the widely used method that allows for the successful mapping of the accessible chromatin regions in the genome. ATAC-Seq relies on the hyperactive Tn5 transposase that fragments the accessible regions in the genome while simultaneously ligating sequencing compatible adaptors (27, 28). Over the years, ATAC-Seq has been applied to many different cell types and tissues (29–31). However, to our knowledge, it has not been performed on a single myofiber, possibly due to the rigidity of their membrane, high levels of mitochondria (32–34) and the low number of myonuclei that is present in a single myofiber (35, 36).

Here, we have adapted OMNI-ATAC-seq to determine genome-wide chromatin accessibility of myonuclei contained within a single Extensor Digitorum Longus (EDL) muscle fiber in mouse. The Single myofiber ATAC-Seq (smfATAC-Seq) method that we applied in this study allows for the investigation of the accessible chromatin state of a single myofiber, without the presence of other confounding cell types, with high sequencing depth. Through this method we provide comparative analysis of chromatin accessibility between resting and regenerating myofibers as well as their MuSC progenitors. This method can be used in the future to profile the epigenetic state of myofibers in diseases and under various physiological and physical stimuli and to identify active cis-regulatory elements in muscle fibers.

## RESULTS

### Generation of ATAC-Seq libraries from a single myofiber

A single EDL myofiber of a mouse contains an average of 200-300 myonuclei (35, 36), making genome-wide analyses of chromatin state difficult. With the advancements in next generation sequencing (NGS) and the development of the OMNI ATAC-Seq protocol (27), analysis of chromatin accessibility of samples with an input of as low as 500 cells is now possible. However, myofibers present additional challenges with their rigid membrane and high levels of mitochondria (32–34). Here we report a robust protocol for the successful application of ATAC-Seq on a single myofiber from the EDL muscle. Our method relies on the lysis and permeabilization of a single myofiber followed by transposition with a hyperactive Tn5 transposase (27) (Figure 1). DNA fragment sizes obtained from the smfATAC-Seq were of a similar range in size as those obtained from conventional OMNI ATAC-Seq that we have performed on MuSCs that were freshly sorted by Fluorescence Activated Cell Sorting (FACS) (Supplemental Figure 1). Furthermore, post sequencing analysis showed that only 0.9-2.09 % of reads were derived from the mitochondria in smfATAC-Seq (Table 1), suggesting that this method is highly efficient for the removal of mitochondria from mitochondria-rich myofibers.

**Figure 1:**
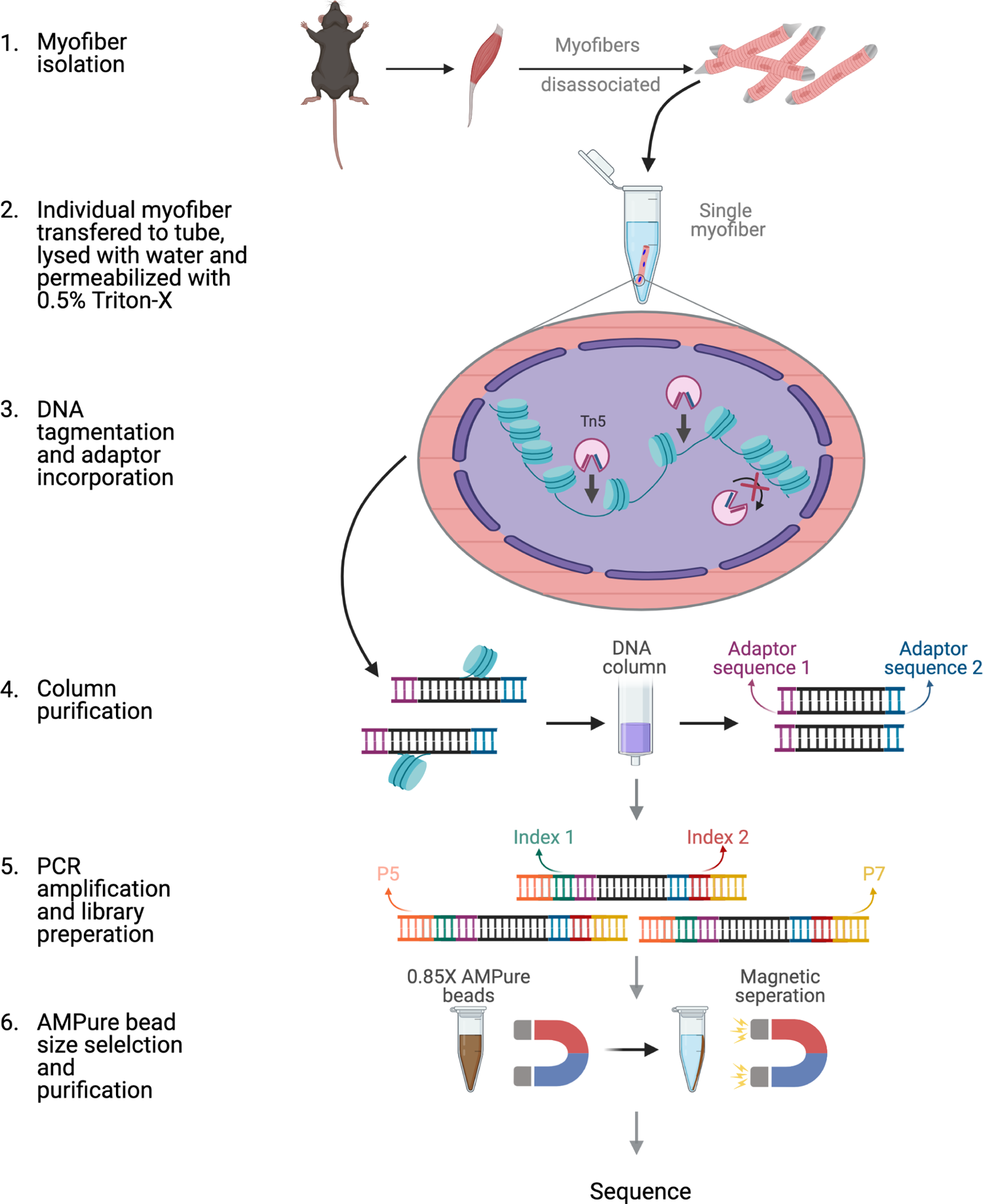
Schematic of ATAC-seq performed on a single myofiber. Schematic showing the steps and reactions involved in the preparation of sequencing ready libraries of single myofiber DNA for ATAC-Seq. Briefly, myofibers were isolated from the EDL muscle and an individual myofiber was transferred to a 0.2mL microtube. The myofiber was then lysed with ddH2O and the myonuclei were permeabilized with 0.5% Triton X-100. Then, open chromatin regions were tagmented with hyperactive Tn5 transposase and the DNA fragments were purified through column purification. The tagmented DNA was then amplified by PCR and Nextera adapters were incorporated. Finally, size selection and purification was performed using 0.85X AMPure beads, resulting in sequencing ready libraries.

**Table 1:** Sequencing read information for smfATAC-Seq and MuSCs ATAC-Seq libraries

| Library | Number of Raw Reads | Number of Surviving Reads | Aligned Filtered Reads (mm10 reference) | Duplicate Reads | Mitochondrial Reads | Percentage of Mitochondrial Reads (%) | Final Reads Aligned | Number of Peaks | Fraction in peaks (FrIP) |
| --- | --- | --- | --- | --- | --- | --- | --- | --- | --- |
| Muscle Stem Cells_1 | 1.76E+08 | 1.14E+08 | 103130186 | 47623836 | 529967 | 0.51 | 5E+07 | 65568 | 0.364 |
| Muscle Stem Cells_2 | 1.75E+08 | 1.17E+08 | 103570009 | 43672484 | 374176 | 0.36 | 6E+07 | 68658 | 0.197 |
| Muscle Stem Cells_3 | 1.32E+08 | 91261584 | 79944121 | 31299456 | 223540 | 0.28 | 5E+07 | 69573 | 0.13 |
| Injured_1 | 2.3E+08 | 1.17E+08 | 90040002 | 81024926 | 830215 | 0.92 | 8E+06 | 32853 | 0.289 |
| Injured_2 | 1.95E+08 | 1.3E+08 | 98752157 | 88549329 | 1300615 | 1.32 | 9E+06 | 28351 | 0.286 |
| Injured_3 | 1.42E+08 | 62888552 | 52132455 | 42271079 | 868808 | 1.67 | 9E+06 | 25002 | 0.233 |
| Uninjured_1 | 1.45E+08 | 75781456 | 61034569 | 52588315 | 1274332 | 2.09 | 7E+06 | 12276 | 0.218 |
| Uninjured_2 | 1.51E+08 | 64192706 | 50120282 | 45914841 | 965037 | 1.93 | 3E+06 | 14742 | 0.321 |

**Table 2:** Percentage of total peaks found in each genomic feature

|  | Muscle Stem Cells (%) | Injured Myofiber (%) | Uninjured Myofiber (%) |
| --- | --- | --- | --- |
| Promoter (<=1 kb) | 20.66 | 31.61 | 56.54 |
| Promoter (1-2 kb) | 4.81 | 4.84 | 3.45 |
| Promoter (2-3 kb) | 4.37 | 3.92 | 3.01 |
| 5'UTR | 0.34 | 0.27 | 0.23 |
| 3'UTR | 2.50 | 1.82 | 1.15 |
| 1 st Exon | 1.83 | 1.47 | 1.53 |
| Other Exon | 4.75 | 3.42 | 2.19 |
| 1 st Intron | 11.85 | 10.87 | 7.35 |
| Other Intron | 20.80 | 18.84 | 10.35 |
| Downstream (<=300 kb) | 1.16 | 1.01 | 0.69 |
| Distal Intergenic | 26.95 | 21.93 | 13.51 |

**Table 3:** Percentage of differential peaks found in each genomic feature

|  | Uninjured Myofiber vs. MuSCs | Uninjured vs Injured Myofiber |
| --- | --- | --- |
| Promoter ($\leq 1$ kb) | 43.07 | 25 |
| Promoter (1-2 kb) | 3.36 | 7.81 |
| Promoter (2-3 kb) | 3.39 | 3.12 |
| 5'UTR | 0.37 | 0.78 |
| 3'UTR | 1.95 | 3.12 |
| 1 st Exon | 2.29 | 3.91 |
| Other Exon | 3.85 | 7.81 |
| 1 st Intron | 9.16 | 13.28 |
| Other Intron | 13.34 | 18.75 |
| Downstream ($\leq 300$ kb) | 0.83 | 0.78 |
| Distal Intergenic | 18.39 | 15.62 |

### smfATAC-Seq can be used to study chromatin accessibility of myofibers under different physiological conditions

In addition to adapting the ATAC-Seq method to study the chromatin accessibility of a single myofiber, we also demonstrate the application of this technique for comparative analysis of chromatin accessibility between myofibers under different conditions. For that purpose, we performed ATAC-Seq on myofibers that were in a resting (uninjured) or regenerating (injured) state. Uninjured and injured (7 days post cardiotoxin (CTX) induced injury) myofibers were isolated from wild type (WT) C57Bl/6 mice and smfATAC-Seq was performed to compare the changes in chromatin accessibility during regeneration. In addition, we compared the chromatin accessibility between myonuclei within a single myofiber and freshly isolated MuSCs. This analysis not only identified accessible regions of chromatin in myofibers and MuSCs but it also revealed a repertoire of active cis-regulatory elements in each sample.

Apart from the myofibers and their associated MuSCs, skeletal muscle also contains many non-myogenic cells such as endothelial cells, adipocytes, hematopoietic cells, fibroblasts, fibro/adipogenic progenitors (FAPs) and macrophages (15,37,38). Our smfATAC-Seq method allows for the analysis of chromatin accessibility of a single myofiber without the confounding effect of these contaminating cell types. To illustrate the absence of the aforementioned cell types in the ATAC-Seq samples, peaks at the promoter regions of marker genes of muscle resident cells were searched for. Specifically, Platelet and Endothelial Cell Adhesion Molecule 1 (*Pecam1*) was used to determine whether endothelial cells were present (39). Similarly, Resistin (*Retn*) and *Cd45* were used as markers for adipocytes and hematopoietic cells, respectively (40, 41). The cell Surface Antigen *Thy1* was the marker selected for fibroblasts (42). In addition, Lymphocyte antigen 6a (*Ly6a*) and Adhesion G protein-coupled receptor E1 (*ADGRE1*) were selected for fibro/adipogenic progenitors (FAPs) and macrophages, respectively (43, 44). None of these marker genes had ATAC-Seq peaks at their promoters, indicating that only a single myofiber is processed without any other contaminating cell types. (Supplemental Figure 2).

smfATAC-Seq may be applied to myofibers under different conditions. For instance, we applied smf-ATAC-Seq to analyze the chromatin accessibility of myofibers under resting and CTX-mediated injury conditions. In a disease condition or in injury, not all myofibers undergo damage or regenerate simultaneously. Therefore, individual myofibers within a muscle can be in different physiological and disease condition asynchronously (45). Damaged or regenerating myofibers can be visualized by their characteristic feature of centrally located nuclei (46). Injured myofibers in this method were visually selected by the presence of the centrally located myonuclei by Hoechst staining and the selected myofiber was used for downstream processing with smfATAC-Seq (Figure 2A, 2B). The selection of a specific myofiber that our smfATAC-Seq allows for, as well as the application of trypsin to remove any confounding resident cell types present, results in the sequencing of DNA fragments corresponding purely to the myonuclei within a specific myofiber.

**Figure 2:**
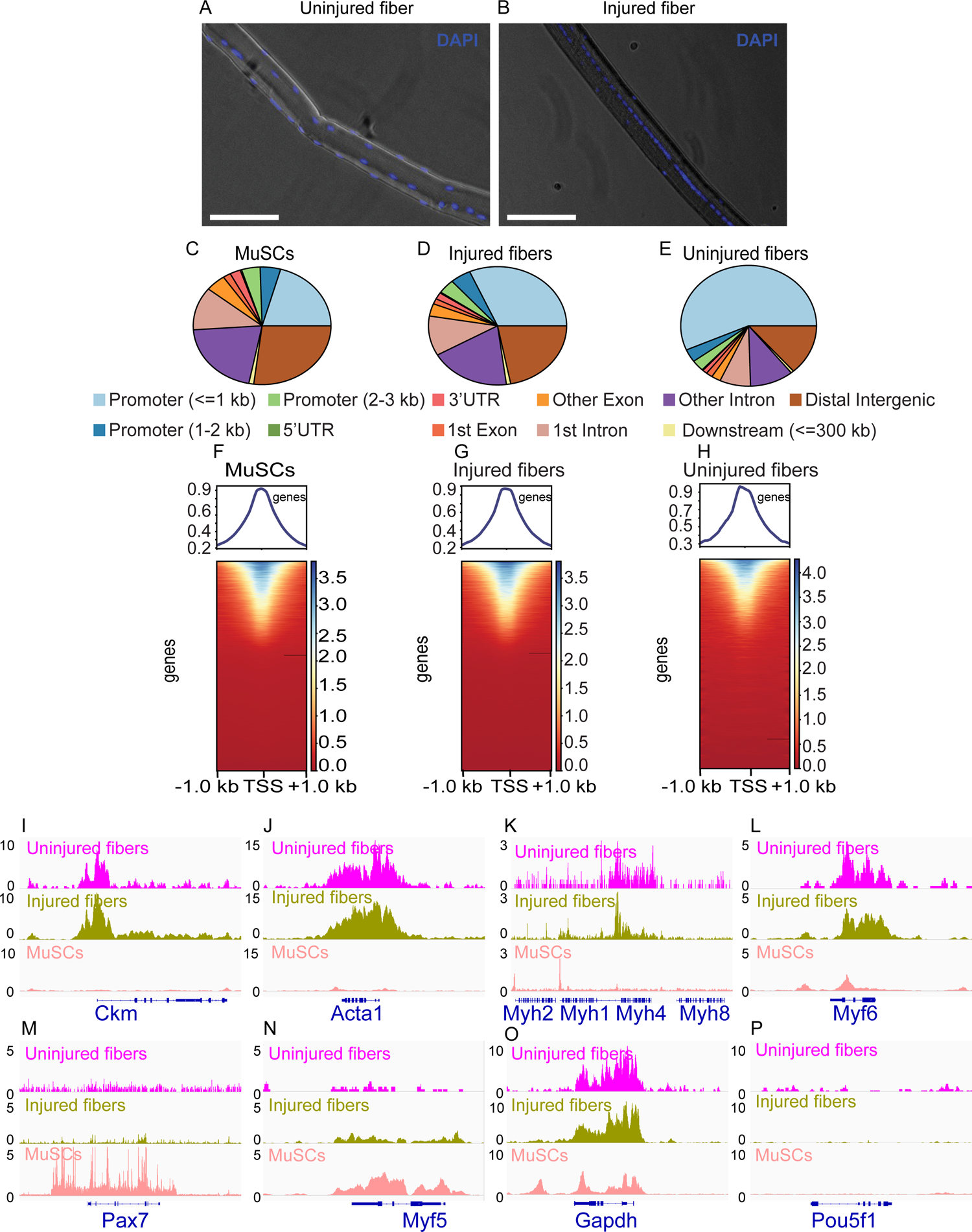
smfATAC-Seq can effectively identify the accessible regions on a single myofiber. (A) Representative picture of an isolated WT C57BL/6J uninjured myofiber stained for DAPI showing presence and location of myonuclei. Scale bar = 50 µm. (B) Representative picture of an isolated WT C57BL/6J injured myofiber (7 days after cardiotoxin induced injury) stained for DAPI showing centrally located myonuclei as a marker of a regenerating fiber. Scale bar = 50 µm. (C-E) Peak annotation pie charts for ATAC-Seq peaks of MuSCs, injured myofibers and uninjured myofibers, respectively. (F-H) Heatmaps showing enrichment at transcription start site (TSS) for the ATAC-Seq libraries of MuSCs, injured myofibers and uninjured myofibers, respectively. (I-P) IGV snapshots of known genes expressed in muscle fiber and/or MuSCs displaying accessibility on their respective TSS. (I) The muscle creatine kinase (*Ckm*). (J) Actin alpha 1 (*Acta1*). (K) Part of the myosin heavy chain (*Myh*) gene cluster. (L) Myogenic factor 6 (*Myf6*). (M) Paired box 7 (*Pax7*). (N) Myogenic factor 5 (*Myf5*). (O) Housekeeping gene *Gapdh*. (P) POU Class 5 homeobox 1 (*Pou5f1*) as a negative control.

### smfATAC-Seq can identify the accessible chromatin regions of a single myofiber

To validate the quality of the ATAC-Seq data generated from a single EDL myofiber, we first investigated the profiles of the ATAC-seq samples from both injured and uninjured myofibers as well as freshly sorted MuSCs. We investigated the similarity between biological replicates for each condition (i.e., uninjured and injured myofibers and MuSCs) by visualization of ATAC-Seq peaks for the muscle-specific gene muscle creatine kinase (*Ckm*) (47), the housekeeping gene *Gapdh* and the MuSC specific gene myogenic factor 5 (*Myf 5*) (48). This analysis not only indicates that smfATAC-Seq can reliably detect chromatin accessibility in a single myofiber but also the presence of comparable peaks in each biological replicate within the specific condition shows the similarity between the samples (Supplemental Figure 3A-C). In addition, Pearson correlation analysis between the biological replicates showed high correlations of ATAC-seq reads between the replicates, indicating consistency within the samples (Supplemental Figure 3D-J). Furthermore, we mapped ATAC-Seq peaks with DNase-Seq from skeletal muscle (Sequence Read Archive, accession # SRX191047) and the similarity between our ATAC-Seq and previous DNase-Seq was confirmed through the similar peaks present for representative genes, as visualized on the IGV (Supplemental Figure 3A-C). These analyses suggest that smfATAC-Seq can robustly measure chromatin accessibility and identify active cis-regulatory elements in a single EDL myofiber.

Following the initial quality control and the correlation analysis, the biological replicates from the same condition were pooled for further analysis. Peak annotation analysis for MuSCs revealed that more than half of the peaks were in the intron/distal intergenic regions (i.e. enhancer regions) and about 25% of the peaks were in the promoter region (Figure 2C, Supplemental Table 2). Peak annotations for the uninjured and injured single myofibers also showed a great proportion of peaks in the enhancer and promoter regions (Figure 2D, 2E, Supplemental Table 2). In addition, an enrichment of ATAC-seq reads around Transcription Start Sites (TSS) from all datasets was observed, which is a typical result expected from ATAC-Seq (30) (Figure 2F-H).

To further assess the quality of the ATAC-Seq data, we analyzed select genes that are expressed by either MuSCs or myofibers. For instance, in the myofiber samples, we confirmed the presence of ATAC-seq peaks in the promoter regions of *Ckm*, Actin alpha 1 (*Acta1*), Myogenic factor 6 (*Myf6*) and Myosin heavy chain 4 (*Myh4*), all of which are expressed by myofibers but not MuSCs (14,47,49,50) (Figure 2I-L). On the other hand, in MuSCs we observed peaks in the promoter regions of Paired box 7 (*Pax7*), and *Myf5,* genes that are known to be expressed in MuSCs (48, 51) (Figure 2M-N). *Gapdh* was used a housekeeping gene for all samples (Figure 2O) and *Pou5f1* was used as a negative control (Figure 2P). These observed peaks for known expressed genes demonstrate that our method, smf-ATAC-Seq can reliably analyse chromatin accessibility in a single myofiber.

### Uninjured and injured myofibers and MuSCs display distinct chromatin states

To show the global differences in chromatin accessibility between MuSCs, uninjured and injured myofibers, we first performed heatmap clustering of Pearson correlation coefficients on all the replicates/samples, which shows that the biological replicates within conditions are more similar to one another than when compared to those from the other conditions (Figure 3A). This can also be observed through Principal Component Analysis (PCA) where each condition clusters separately, with the injured and uninjured myofibers being more similar to one another than to MuSCs (Figure 3B).

**Figure 3:**
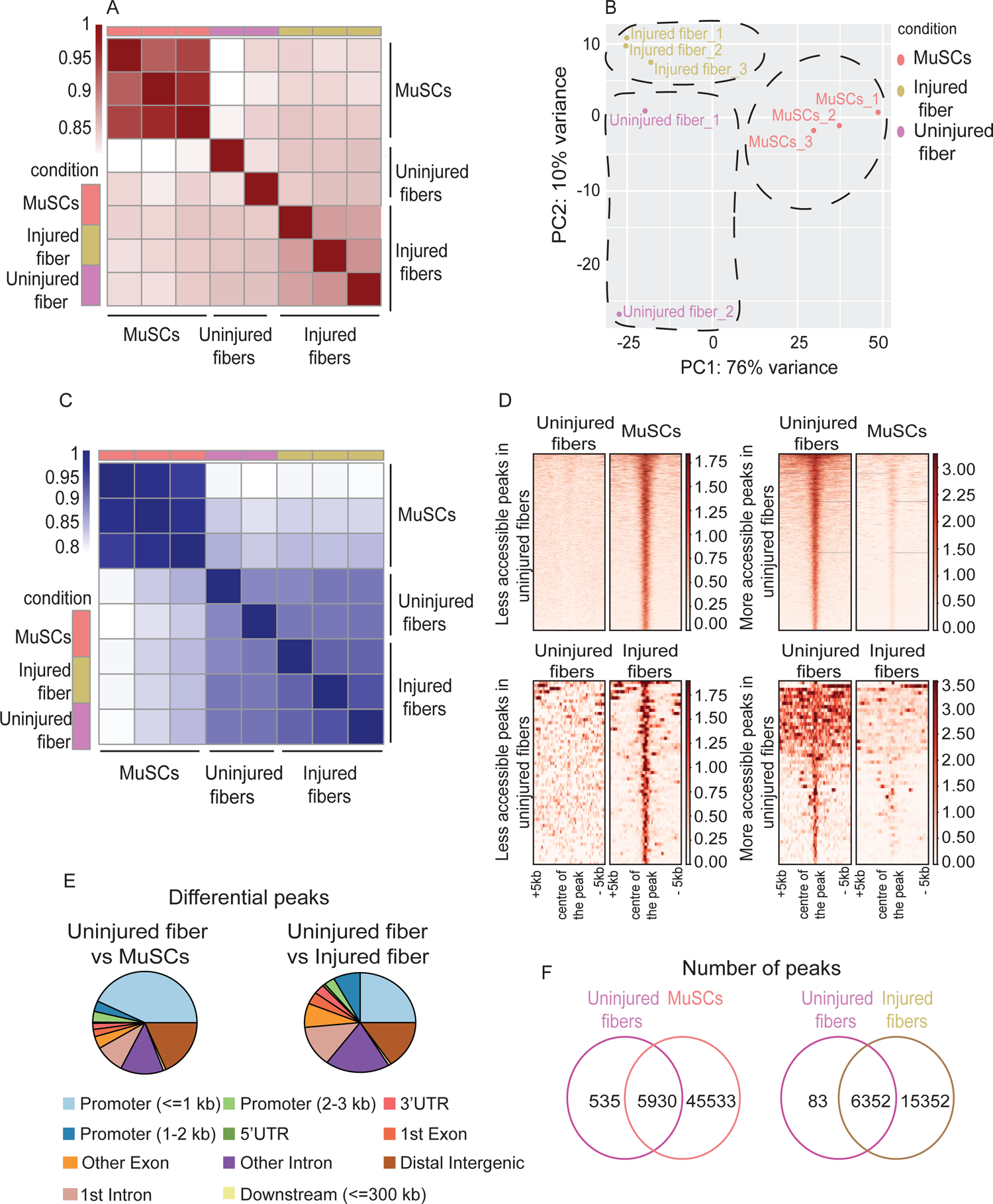
Uninjured and injured myofibers and MuSCs display distinct chromatin states. (A)Heatmap clustering of Pearson correlation coefficients showing the correlation between the replicates of the conditions in the regions defined by the union peakset (merged peaks of all replicates/samples). (B) Projection of samples along first two principal components found by PCA showing the separate clustering of different samples and the clustering of each replicate of the same condition together. (C) Pile up analysis of differentially accessible peaks between uninjured myofibers and MuSCs and between injured myofibers and uninjured myofibers. Less accessible regions: FDR < 0.05 and LFC < 0.5. More accessible peaks: FDR < 0.05 and LFC >2. (D) Heatmap clustering of Pearson correlation coefficients indicating the correlation between the replicates in the regions defined by the consensus peakset derived from the uninjured myofibers, injured myofibers and MuSCs. (E) Peak annotation pie charts for the differentially accessible peaks between uninjured myofibers vs MuSCs and uninjured myofibers vs injured myofibers. (F) Venn diagram of the number of ATAC-Seq peaks that are unique or overlapping between uninjured myofibers vs MuSCs and uninjured myofibers vs injured myofibers.

To ensure that the differences seen between myofibers were not due to differences in fiber types, we investigated the chromatin accessibility of known marker genes for slow and fast fiber types. Troponin I2 (*Tnni2*) and Troponin T3 (*Tnnt3*), markers of fast fiber types, (52, 53) had a high level of chromatin accessibility while Troponin T1 (*Tnnt1*) and Myosin heavy chain 7 (*Myh7*), expressed in slow fiber types showed no chromatin accessibility (53, 54) (Supplemental Figure 4). This data indicates that only fast fiber types were analyzed in this study and that the differences in the chromatin state between the injured and uninjured myofibers were not due to the differences in the fiber types.

For further comparison of accessible regions between the conditions, we performed differential analysis of ATAC-Seq peaks based on the regions defined by the consensus peak sets derived from the uninjured and injured myofibers and MuSCs conditions. The clustering analysis based on the consensus peak set shows the overall chromatin state differences between the MuSCs and injured and unjured myofibers where the replicates of a condition are more similar to one another than to other conditions (Figure 3C).

The unique chromatin state in each condition can be can be observed through the pile up analysis that we have performed for the more accessible peaks (LFC> 2) and less accessible peaks (LFC < 0.5) between uninjured myofibers and MuSCs as well as between uninjured and injured myofibers (Figure 3D). In addition, the proportion of the differential peaks corresponding to various genomic regions such as promoters and enhancers, differs depending on whether we compare MuSCs to myofibers or compare the myofibers during regeneration and homeostasis. For instance, the differential peaks between uninjured myofibers and MuSCs were mostly found close to the promoter region (≤ 1 kb) whereas in the uninjured and injured myofibers comparison, a greater proportion of differential peaks were found in the intron/distal intergenic regions (i.e. enhancer regions) (Figure 3E). This implies that MuSCs and myofibers mostly differ in their promoter accessibility, whereas myofibers during homeostasis and regeneration differ mostly at the level of distal regulatory elements.

Furthermore, we performed occupancy analysis (using DiffBind) in order to determine the unique and common peaks for each comparison between the conditions. Since the occupancy analysis relies on the peak score, the distribution pattern of the peak scores for all the conditions was assessed and found similar despite the differences in the total number of peaks between the conditions (Supplemental Figure 5A). Occupancy analysis between uninjured myofibers and MuSCs revealed that MuSCs have 45533 unique peaks while myofibers contain only 535 unique peaks which are not present in MuSCs (Figure 3F). There are also many common accessible regions as seen from 5930 peaks that are common to both uninjured myofibers and MuSCs. This analysis suggests that myonuclei share large number of open chromatin with their parental stem cells. On the other hand, the occupancy analysis between uninjured and injured myofibers revealed that there are 6352 overlapping peaks between the regenerating and resting myofibers. However, this analysis also revealed that there are 15352 unique peaks in the injured myofibers and only 83 peaks that are unique to the resting myofiber (Figure 3F). Furthermore, when comparing the read count between conditions around the center of unique peaks, it can be observed that each condition displays a unique open chromatin signature (Supplemental Figure 5B, 5C).

### Comparative analysis of the chromatin state between MuSCs and myofibers

To get a better understanding of the functional differences in chromatin accessibility between MuSCs and myofibers, we first performed Gene Ontology (GO Biological Process) analysis on the genes associated with the nearest peaks from the uninjured myofiber and on the genes associated with the nearest unique peaks in the myofiber compared to MuSCs. As expected, this revealed myofiber specific biological processes such as myofiber structure and organization (Figure 4A and Supplemental Figure 6A). On the other hand, GO term analysis on the genes associated with the nearest peaks from MuSCs as well as the genes associated with the nearest unique peaks to MuSCs revealed biological processes such as adherens junction organization, membrane permeability and regulation of notch signaling which play key roles in MuSCs quiescence and function (55, 56) (Figure 4B and Supplemental Figure 6B). The analysis above also revealed that genomic regions that remain in an open chromatin state when muscle stem cells fully differentiate into myofibers correspond to genes that are involved in processes such as mitochondrial transport, regulation of transcription and regulation of metabolites and energy (Supplemental Figure 6C). The changes in the chromatin state between MuSCs and myofibers can also be observed from the volcano plots showing differential peaks between conditions labeled by their nearest gene (Figure 4D). For instance, genomic regions associated with genes such as muscle-specific titin-capping protein (*Tcap*), a component of the skeletal muscle z-disc, as well as genes that are involved in regulatory and structural functions in skeletal muscle such as titin gene (*Ttn*) are associated with more accessible chromatin regions in the myofiber compared to MuSCs (57, 58) (Figure 4D).

**Figure 4:**
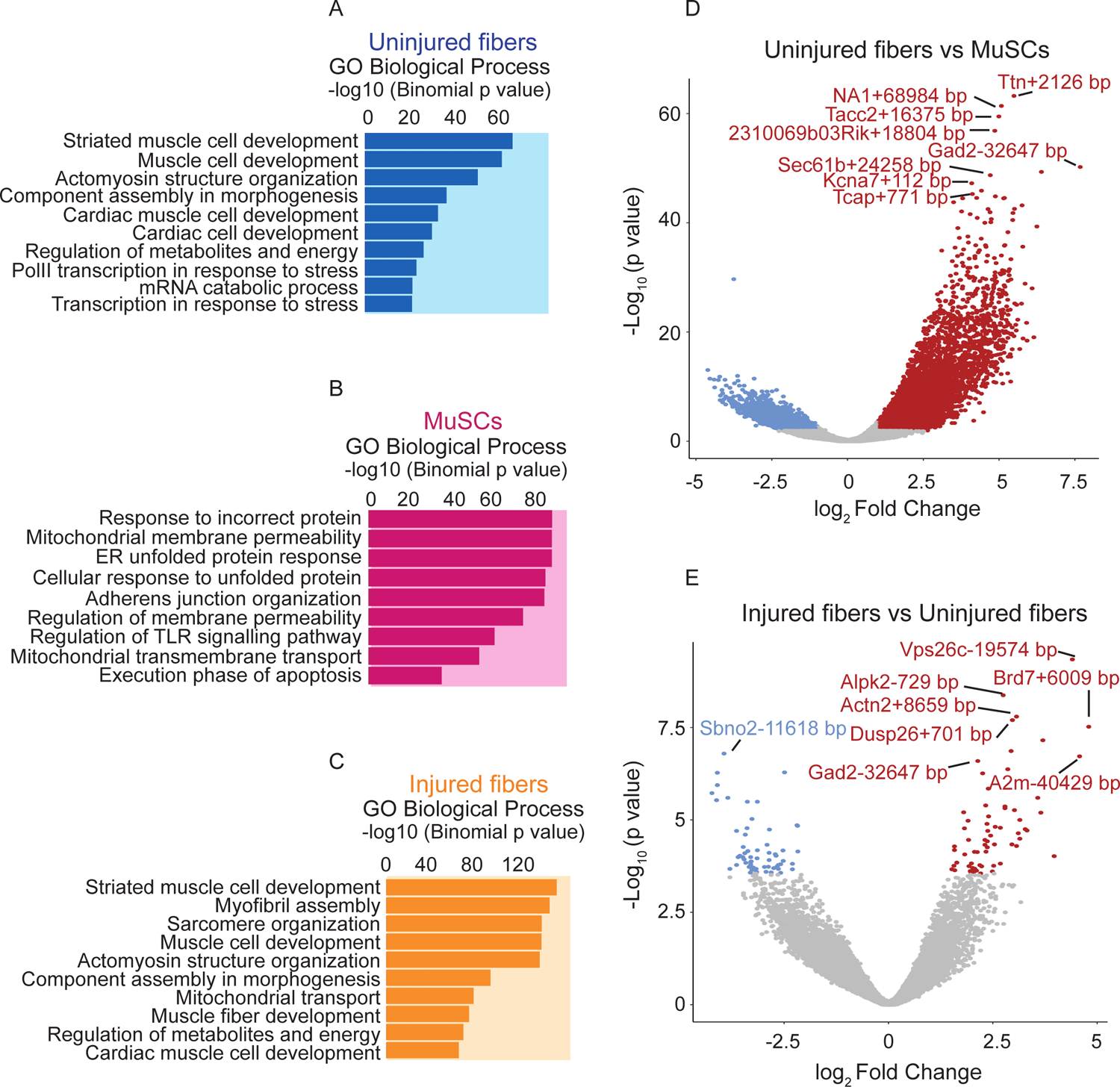
Comparative analysis of chromatin state between uninjured myofibers and MuSCs and between uninjured myofibers and injured myofibers. (A-C) Gene Ontology (GO Biological Process) analysis of genes associated with ATAC-Seq peaks based on association by proximity using Genomic Regions Enrichment of Annotations Tool (GREAT) (84) for all peaks present in the uninjured myofibers, MuSCs and injured myofibers respectively. (D) Volcano plot of differentially accessible regions/peaks identified by FDR < 0.05 and LFC >= 1 between uninjured myofibers and MuSCs. Each dot represents a differentially accessible region/peak and the distance to the nearest gene is annotated. (E) Volcano plot of differentially accessible regions/peaks identified by FDR < 0.05 and LFC >= 1 between uninjured myofibers and injured myofibers. Each coloured dot represents a differentially accessible region/peak and the distance to the nearest gene is annotated.

Additionally, we performed GO term analysis between uninjured and injured myofibers, which revealed that the accessible regions in the resting and regenerating myofibers corresponded to genes involved in similar processes. GO term analysis on the genes associated with the nearest peaks from uninjured myofibers and from the injured myofibers as well as the genes associated with the nearest common peaks between uninjured and injured myofibers revealed biological processes involved in striated muscle cell development, actomyosin structure and sarcomere organization, which are important for myofiber structural formation and for the proper function of myofibers (Figure 4A, 4C, Supplemental Figure 6D). On the other hand, genes associated with the nearest unique peaks from the injured myofibers mostly belong to processes involved in structural components of the myofiber while the genes associated with the nearest unique peaks from uninjured myofibers correspond to genes involved in ion transport and metabolism (Supplemental Figure 6E-F).

### Identification of cell type specific pathways by global analysis of chromatin accessibility

To further understand the functional differences in chromatin accessibility between different cell types, we investigated the cell-type-specific pathways. To accomplish this, Gene Set Enrichment Analysis (GSEA) was performed on genes associated with differentially accessible peaks between the conditions. Importantly, the GSEA between uninjured and injured myofibers revealed that inflammatory response and Il2-Stat5 signaling and injury related pathways are still operational even after seven days of CTX-mediated injury to muscle (59) (Figure 5A).

**Figure 5:**
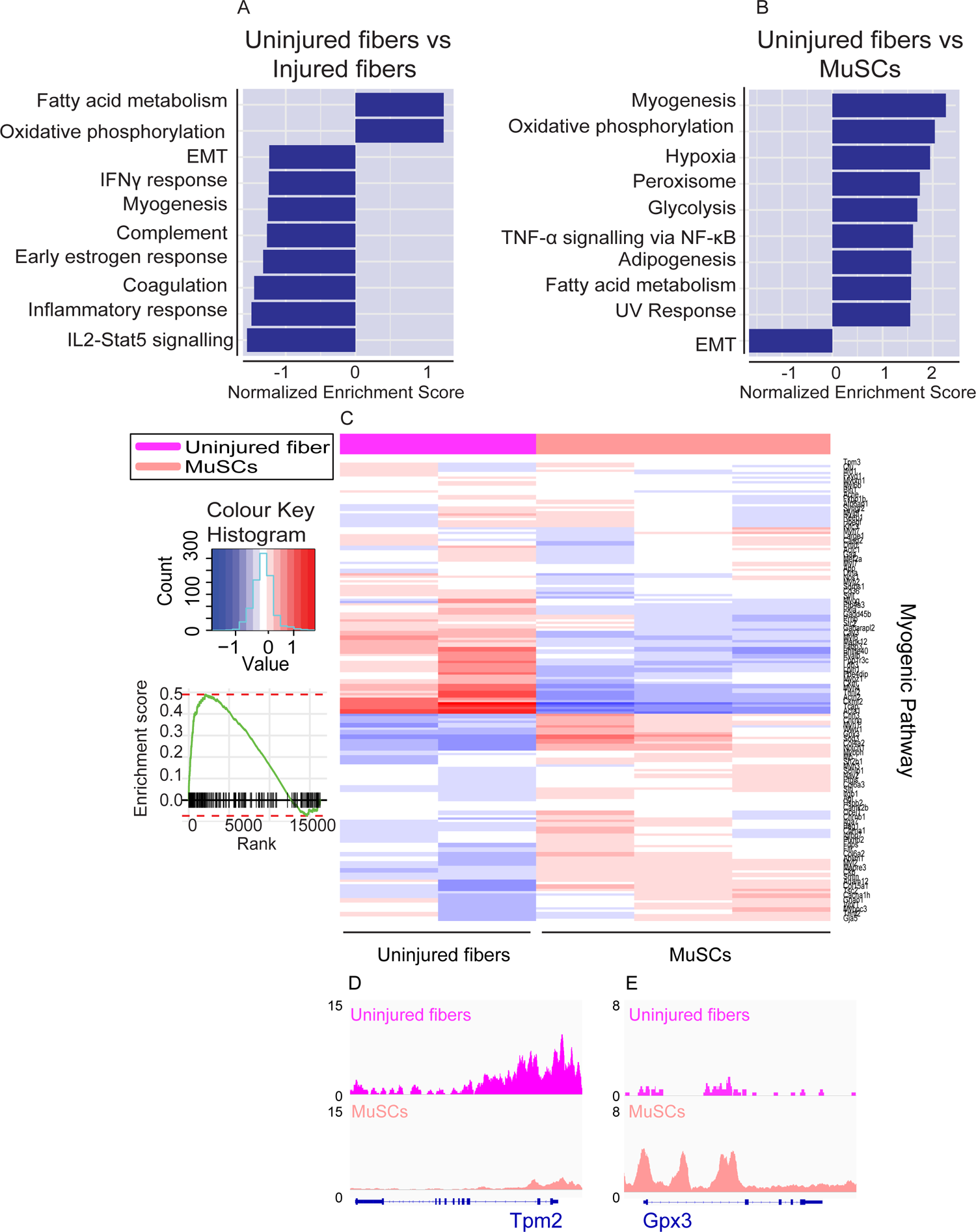
Identification of cell type specific pathways by global analysis of chromatin accessibility. (A) Gene Set Enrichment Analysis performed on nearby genes of differentially accessible regions/peaks for uninjured myofibers compared to injured myofibers. Top 10 enriched pathways are shown although do not reach significance. (B) Gene Set Enrichment Analysis performed on nearby genes of differentially accessible regions/peaks for uninjured fibers compared to MuSCs. Top 10 significantly enriched pathways are shown (FDR < 0.01). (C) Heatmap for myogenesis based on read counts of MuSCs and uninjured fibers +/- 1 kb of the TSS of each gene in the myogenic pathway. (D) IGV snapshot of Tropomyosin 2 (*Tpm2*). (E) IGV snapshot of Glutathione Peroxidase 3 (*Gpx3*).

The GSEA between uninjured myofibers and MuSCs revealed that one of the top significantly enriched pathways is myogenesis where we can observe that genes associated with differentiation and myofiber function such as *Myf6*, *Ckm* and Tropomyosin 2 (*Tpm2*) (14,47,60,61) have higher accessibility in the myofiber compared to MuSCs (Figure 5B-5D). On the other hand, genes associated with quiescence and MuSCs such as alpha 7 integrin (*Itga7*) and *Gpx3* are more accessible in MuSCs compared to the myofiber, as expected (62, 63) (Figure 5C, 5E).

Moreover, differential chromatin accessibility between the MuSCs and their myofiber derivatives show differences in pathways that are known to be important for muscle, such as Notch and TGFβ signalling (55,56,64,65) (Supplemental Figure 7). For example, increased accessibility of *Notch1* is seen in MuSCs while increased accessibility of Jagged-2 (*Jag2*) is observed in the myofibers regardless of whether they are regenerating or homeostatic as seen by the height of the peaks at their promoters (Supplemental Figure 7A-C). On the other hand, for TGFβ signalling, Noggin (*Nog*) shows more accessibility in the myofibers while bone morphogenetic protein-4 (*Bmp4*) has increased accessibility in MuSCs (Supplemental Figure 7D-F). Taken together, this data shows that smfATAC-Seq is an effective method to analyze chromatin accessibility and to identify active cis-regulatory elements in a single muscle fiber as well as to compare muscle fibers under different physiological conditions.

## DISCUSSION

Analysis of the myofiber-specific chromatin state and gene expression profile is very limited due to the heterogenous nature of muscle with the presence of numerous non-myogenic cells in the tissue. Whole muscle or muscle biopsies represents a pooled result of numerous cell types which are present in the muscle tissue. To overcome this limitation, single nucleus RNA-Seq (snRNA-Seq) and ATAC-Seq (snATAC-Seq) have been developed and performed on myonuclei to allow for the computational removal of other cell types (16–18). However, these methods still sequence all myonuclei present in the muscle and cannot distinguish between different myofibers within a muscle. Although snATAC-Seq provides high dimensionality, it is limited in sequencing depth due to the generation of sparse reads. Although computational pseudo bulking of snATAC-Seq can increase read numbers for comparative analysis between samples and conditions, the pooled reads represent average of all myonuclei within the sample.

In this study we have introduced a highly effective protocol based on adaption of OMNI ATAC-Seq (27) to quantify chromatin accessibility of a single EDL myofiber with high resolution and sequencing depth. This method allows comparative analysis of chromatin accessibility within and between muscle types with a potential wide-spread use in future studies to investigate fiber-specific epigenetic alterations in skeletal muscle.

The smfATAC-Seq protocol that we introduce in this study investigates the open chromatin state of myofibers at a single myofiber resolution, and with a high sequencing depth that allows for peak calling and differential peak analysis. Using this method, we have demonstrated that accessible chromatin regions of myonuclei contained within a single EDL myofiber can be tagmented and that high-quality sequencing ready libraries can be generated from these fragments. Sequencing of these libraries allow for sufficient depth and peak calling that can be used for genome-wide analysis of chromatin accessibility between myofibers. In this study, we have also demonstrated that smfATAC-Seq strictly investigates a single myofiber without the confounding presence of muscle resident, non-myogenic cell types. Additionally, application of trypsin to isolated myofibers effectively removes MuSCs that are associated with myofibers (66) and was confirmed by the absence of peaks at the promoters of known genes associated with muscle stem and niche cells. Although all of the smfATAC-Seq samples sequenced in this study were fast type myofibers, this protocol can be used to distinguish between different fiber types which could be applied to study myofiber heterogeneity under various physiological conditions. smfATAC-Seq peaks associated with genes involved in muscle structure and function such as *Acta1*, *Ckm* and the myosin heavy chain cluster, indicates that our smfATAC-Seq is a robust technique to investigate genome-wide chromatin accessibility of a single myofiber.

A key implication of this technique is its applicability to the study of changes in chromatin accessibility between myofibers in different contexts. As a demonstration of this, we have performed smfATAC-Seq on uninjured myofibers as well as injured myofibers isolated seven days post-injury to investigate the changes in chromatin accessibility that occurs during regeneration. Through Pearson correlations and PCA analysis, we showed that each cell type including resting and injured myofibers cluster separately, indicating the power of smfATAC-Seq to determine chromatin signature from minute materials such as a single myofiber. In addition, through occupancy analysis, we showed that there is a large difference in the number of unique peaks present in the injured myofibers compared to uninjured myofibers. This indicates that there are major modifications to chromatin accessibility in the context of regeneration. However, GO term analysis of genes associated with the accessible chromatin regions in both injured and uninjured fibers display significant similarities in the biological processes and pathways such as striated muscle cell development, actomyosin structure and sarcomere organization, which are key factors for the proper structure and function of muscle. Despite these similarities, there are certain trends in which uninjured myofibers have increased accessibility in genes involved in energy metabolism, while injured myofibers have greater accessibility in genes involved in myogenesis and inflammatory response which is what would be expected in the case of an injury and regeneration (67). Despite the increase in chromatin accessibility during injury, the accessible chromatin regions in both injured and uninjured fibers are associated with genes involved in similar biological processes. This similarity at the gene network despite differences in chromatin profile may suggest activation of multiple enhancers on core muscle structural genes in the case of injury. Another possible reason could be the length of the recovery time where at seven days post injury, a significant portion of early activated genes may have gone back to steady state. Previously, a study investigating the changes in the transcriptional profile of MuSCs and various muscle resident cells throughout different time points of muscle injury using single cell RNA-Seq, revealed that after seven days of regeneration most cell types returned to a state that was similar to homeostasis (37). Therefore, it is possible that harvesting the injured EDL myofibers seven days post injury allowed these myofibers to return to a state reminiscent of homeostatic myofibers. smfATAC-Seq can be further implemented to study the effects of regeneration on chromatin accessibility at different time points. Further, our analyses of the myofibers in this study indicates that this technique can effectively compare samples between conditions and could see future use in the study of chromatin accessibility of fibers under different biologically relevant conditions.

Lastly, we have also used smfATAC-Seq to compare changes in chromatin accessibility between MuSCs and myofibers. Our data shows that the regions of open chromatin in the myofibers correspond to genes involved in structural components of the muscle, such as the z-disc, which are important for the proper functioning of the muscle. On the other hand, open regions of chromtain in the MuSCs mostly correspond to genes involved in membrane permeability, adherens junction organization and signalling pathways implicated in the regulation of MuSC function (68). The analysis also revealed that chromatin regions that are accessible in both MuSCs and myofibers correspond to genes that are crucial for general function of cells such as those involved in mitochondrial transport, regulation of transcription and regulation of metabolites and energy.

Overall, smfATAC-Seq is a robust molecular tool that can be used to analyze genome-wide chromatin accessibility of a single myofiber. The sequencing depth from this approach, allows for in depth analysis, peak calling, quantitative analysis of chromatin accessibility and to identify active enhancers and promoters in a single muscle fiber. smfATAC-Seq can be used to study epigenetic alterations that occur in muscle fibers during development, and in response to exercise.

## MATERIALS AND METHODS

### ATAC-Seq on a Single Myofiber

#### Isolation of Extensor Digitorum Longus (EDL) from cardiotoxin-induced injured muscle

The Extensor Digitorum Longus (EDL) muscle was injured by intramuscular injection of 50 µL of 5 µM cardiotoxin (CTX) (Sigma, 11061-96-4). Mice were treated with carprofen 20 minutes prior to CTX injection and were injected with CTX under anesthesia by isoflurane. Mice were sacrificed seven days post injury and the EDL was collected from the hind limb of each mouse with the contra lateral EDL being used for the isolation of uninjured myofibers.

#### Dissection of EDL muscle

The EDL muscle was dissected as previously described (66). Briefly, the skin of the hindlimb was removed and the tibialis anterior (TA) muscle was excised with a pair of dissection scissors. The tendons of the EDL were exposed and cut from tendon to tendon with scissors.

#### Isolation of a single EDL myofiber

Individual myofibers were isolated from the EDL muscle as previously described (66). Briefly, the intact EDL muscle was placed in a 1.5 mL eppendorf tube with 800 µL of myofiber digestion buffer containing 1000 U/mL of collagenase from *Clostridium histolyticum* (Sigma, C0130) in un-supplemented DMEM (Gibco, 11995-065) for 1 hour. Trypsin was added to the myofiber digestion buffer at a final concentration of 0.25% to remove the myofiber associated muscle stem cells. The EDL myofibers were then transferred into 2 mL of 1X PBS (Wisent, 311-425-CL) in a 6 well-plate that had previously been coated with DMEM supplemented with 10% horse serum (HS) (Wisent, 065250). The EDL was then gently pipetted up and down with a large-bore glass pipette to disassociate the myofibers.

#### Selection of injured and uninjured myofibers

Live myofibers in the 6-well plate were stained with 2 µL of 5mg/mL of Hoechst (Molecular Probes, H1399) in 2 mL of 1X PBS for 5 minutes in a 37 °C with 5% CO2 incubator. The myofibers were then visualized under a microscope and selected based on the myonuclei location, where myofibers with a pattern of centrally located nuclei were determined to be regenerating and picked for the injury condition. Individual myofibers were then transferred to 0.2 mL microtubes using a small-bore glass pipette coated with HS.

#### Lysis and permeabilization of the myofiber

Residual media was removed with pipette under a microscope. Individual myofibers in 0.2 mL microtubes were put in 10 µL of ddH2O for 5 minutes on ice. The ddH2O was removed with a pipette under a microscope, ensuring that the myofiber remained in the tube. The myofiber was then permeabilized with 20 µL of 0.5% Triton X-100 (Sigma, T9284) in PBS for 15 minutes at room temperature (RT). The permeabilization buffer was removed with a pipette under a microscope and the myofiber was washed twice with 200 µL of 1X PBS.

#### Tagmentation of the myofiber by Tn5 transposase

Transposition and ATAC-seq library preparation for a single myofiber was adapted from previously described OMNI ATAC-Seq protocol (27). The permeabilized myonuclei were tagmented with tagmentation mixture optimized for use on a myofiber (20 µL Tagment DNA Buffer (TD Buffer) (Illumina, 20034197), 13.3 µL PBS, 0.2% Tween-20 (Sigma, P1379-1L), 0.02% Digitonin (Promega, G9441), 1.39 µL Tn5 (Illumina, 20034197) and 4.61 µL water). Each single myofiber was incubated with 6 µL of the tagmentation mixture at 37 °C for 56 minutes with periodic shaking of the tubes every 5-7 minutes. Following the transposition with Tn5, DNA was purified using a QIAquick PCR Purification Kit (Qiagen, 28104) according to the manufacturer’s guidelines.

#### Library preparation

The purified DNA was PCR amplified for 15 cycles using Q5 High Fidelity DNA polymerase (New England Biolabs, M0491S) with the incorporation of Illumina Nextera XT adaptors (Illumina, FC-131-1001). The libraries were then size selected with AmpureXP Beads (Beckman, Cat# A63880) at a 1: 0.85 ratio (v/v). The size selected libraries were verified for quality control by bioanalyzer as well as verification of the library size via visualization on an agarose gel stained with GelGreen dye (Biotium, 41005). Libraries were then sequenced on NovaSeq6000 Sprime Paired End (PE) 150 bp.

### ATAC-Seq on MuSCs

#### Isolation of MuSCs by fluorescence-activated cell sorting (FACS) for ATAC-Seq

MuSCs were isolated by Fluorescence Activated Cell Sorting (FACS) as previously described (69). Briefly, hindlimb muscles from Pax7/GFP^+^ mice were dissected and chopped. The minced muscles were then transferred into a 15 mL Falcon tube and digested in un-supplemented F10 media (Gibco, 11550043) with 2.4 U/mL Collagenase D (Roche, 11088882001), 12 U/mL Dispase II (Roche, 39307800) and 0.5mM CaCl_2_. Digestion was performed on a shaker in an incubator at 37 °C with 5% CO_2_ for 30 minutes. Following the first digestion, digested muscles were centrifuged at 600 g for 20 seconds and the supernatant was transferred to a 50 mL Falcon tube with 9 mL FBS (Wisent, 080450) and was kept on ice. The remaining pellet was triturated and was digested for another 15 minutes with additional digestion buffer added. After the final digestion, the digested muscle mixture was transferred to the 50 ml Falcon tube containing the previously digested mixture. The digested muscle mixture was then filtered through a 40 µm cell strainer (Falcon, C352340) and was centrifuged at 600 g for 18 minutes at 4°C. The pelleted cells were then resuspended in 800 μL FACS buffer that is composed of 2% FBS/ PBS (v:v), 0.5 mM EDTA (Invitrogen, AM9261) and with 0.5 μL DAPI (5mg/mL) (Invitrogen, D3671). Resuspended cells were then filtered through 40 µm cell strainer and were transferred into polypropylene round-bottom FACS compatible tubes (Falcon, 352063). MuSCs were sorted with a FACSAria Fusion cytometer (BD Biosciences) based on negative selection for DAPI and positive selection for GFP.

#### Lysis and transposition of MuSCs

ATAC-Seq on MuSCs was performed based on the previously established OMNI-ATAC-Seq protocol (27). Briefly, five thousand MuSCs were sorted by FACS into 30 μL of the ATAC lysis buffer containing 10 mM Tris-HCl (pH 7.5), 10 mM NaCl (Bioshop, 7647-14-5), 3 mM MgCl2 (Sigma, 7786-30-3), 0.1% Tween-20 (Sigma, P1379-1L), 0.1% NP-40 (Sigma, 74385), and 0.01% Digitonin (Promega, G9441) in a 0.2 mL microtube. Cells were incubated in the lysis buffer for 5 minutes on ice and then 3 minutes at room temperature (RT). Cells were then washed with 100 µL of wash buffer composed of 10mM Tris-HCl (pH 7.5), 10Mm NaCl, 3mM MgCl2 and 0.1% Tween-20, and were centrifuged at 800 g for 10 minutes. The pellet was resuspended in 10 µL of transposition mixture (5 µL TD buffer, 3.2 µL PBS, 0.89 µL Tn5 (Illumina, 20034197), 0.1% Tween-20, 0.01% Digitonin and 0.75 µL nuclease free water). Transposition was performed for 20 minutes at 37 °C while shaking the tubes every 5-7 minutes. The DNA was then purified using a QIAquick PCR Purification Kit according to the manufacturer’s guidelines.

#### Library preparation for MuSCs ATAC-Seq

The eluted tagmented DNA was PCR amplified for 12 cycles with the incorporation of Illumina Nextera XT adapters using Q5 High Fidelity DNA polymerase. The libraries were then size selected with AmpureXP Beads at a 1: 0.85 ratio (v/v). The libraries were then verified by bioanalyzer and agarose gel visualization. Finally, the samples were sequenced on NovaSeq6000 Sprime Paired End (PE) 150 bp.

#### ATAC-Seq data processing

The sequencing data was processed using the GenPipes pipeline v.3.1.5 (70). The raw reads were trimmed using Trimmomatic v.0.36 (71) and aligned to the mm10 genome assembly using the Burrows-Wheeler Aligner v.0.7.12 (72). Reads were filtered to keep only high quality alignments (MAPQ score > 20) and duplicates were removed using SAMtools v.1.3.1 (73). Peak calling was performed with MACS2 v.2.1.1 (74) using piling up of paired-end fragment mode (*--format BAMPE*). The peak files (bed) were filtered by removing the ENCODE black listed regions (https://www.encodeproject.org/files/ENCFF547MET) using BEDTools v2.29.1 (75). Mitochondrial reads were also removed before the analysis.

#### Correlation analysis between the biological replicates and clustering

In order to perform a quantitative comparison of the read counts within accessible regions, the overlapping peaks of all replicates were merged using BEDTools v2.29.1 (75). This set of merged peaks and the BAM alignment files were used as input for the *featureCounts* function of Rsubread v.2.2.6 (76) to generate a raw-count matrix. The raw counts were normalized by rlog transformation using DESeq2 (77) with respect to library size. Pearson correlation coefficients were calculated based on the normalized counts for each pairwise comparison. Principal component analysis (PCA) and hierarchical clustering were also performed to evaluate the similarity between the replicates.

#### Peak annotation analysis

For each condition, the BAM alignment files of the replicates were merged and peak calling was performed with MACS2 v.2.1.1 (74). Peak sets for each condition were annotated using the ChIPseaker v.1.24.0 (78) *annotatePeak* function, and the UCSC Genome Browser knownGene (mm10) table.

#### Obtaining coverage tracks

The BAM alignment files were converted to bigWig format and normalized by scaling factor (-- scaleFactor) with the deepTools v.2.5.0.1 (79) *bamCoverage* function.

#### Enrichment of genomic signal around TSS

The bigWig files and the TSS coordinates obtained from the UCSC Genome Browser knownGene (mm10) table were used as input for the *computeMatrix* function of deepTools v.2.5.0.1 (79). This matrix was used for *plotHeatmap* function to generate the heatmap.

#### Identification of overlapping/unique accessible regions

For each comparison between the conditions, overlapping and unique accessible regions were identified with DiffBind v.2.16.2(80) based on the measure of confidence in the peak call by MACS2 v.2.1.1 (74).

#### Analysis of differentially accessible regions

The identification of differentially accessible regions (DARs) between the conditions was done using DiffBind v.2.16.2 (80) and edgeR v.3.30.1 (81). Log fold changes were calculated, and their associated p-values were corrected for multiple hypothesis testing via the Benjamini–Hochberg procedure to obtain adjusted p-values. The DARs were annotated by their nearest gene using the *annotatePeaks.pl* function of Homer v.4.11(82).

#### Gene set enrichment analysis

Genes nearby the DARs were ranked based on the log-fold change calculated with edgeR v.3.30.1 (81). This ranked list of genes was used as input to perform gene set enrichment analysis with the *fgseaMultilevel* function of the R package fgsea v.1.14.0 (83). The *FGSEA-multilevel* method is based on an adaptive multi-level split Monte Carlo scheme, which allows the estimation of very low p-values. The Hallmark gene sets collection from the Molecular Signatures Database (MSigDB) (83) was used as a reference to identify the biological processes that were significantly enriched.

#### Animal care

All procedures that were performed on animals were approved by the McGill University Animal Care Committee (UACC).

## Data availability

The data discussed in this study have been deposited in NCBI’s Gene Expression Omnibus and are accessible through GEO Series accession numbers GSE173676 and GSE171534.

## Author contributions

Conceptualization: VDS; Methodology: KS, DMB, VDS; Investigation: VDS, MMS, KS, DMB, TK; Visualization: KS, VDS, MMS, DMB; Funding acquisition: VDS; Project administration: VDS; Supervision: VDS, GB; Software: MMS; Writing – original draft: KS, DMB; Writing – review & editing: KS, DMB, VDS, MMS, FL.

## Acknowledgments

We thank Christian Young at the Lady Davis Institute for Medical Research—Jewish General Hospital-core facility for his help with fluorescence-activated cell sorting (FACS) of muscle stem cells. We thank Dr. Michael Witcher at McGill University Department of Oncology for his critical comments and careful review of an early draft of this manuscript.

---

**Figure S1:**
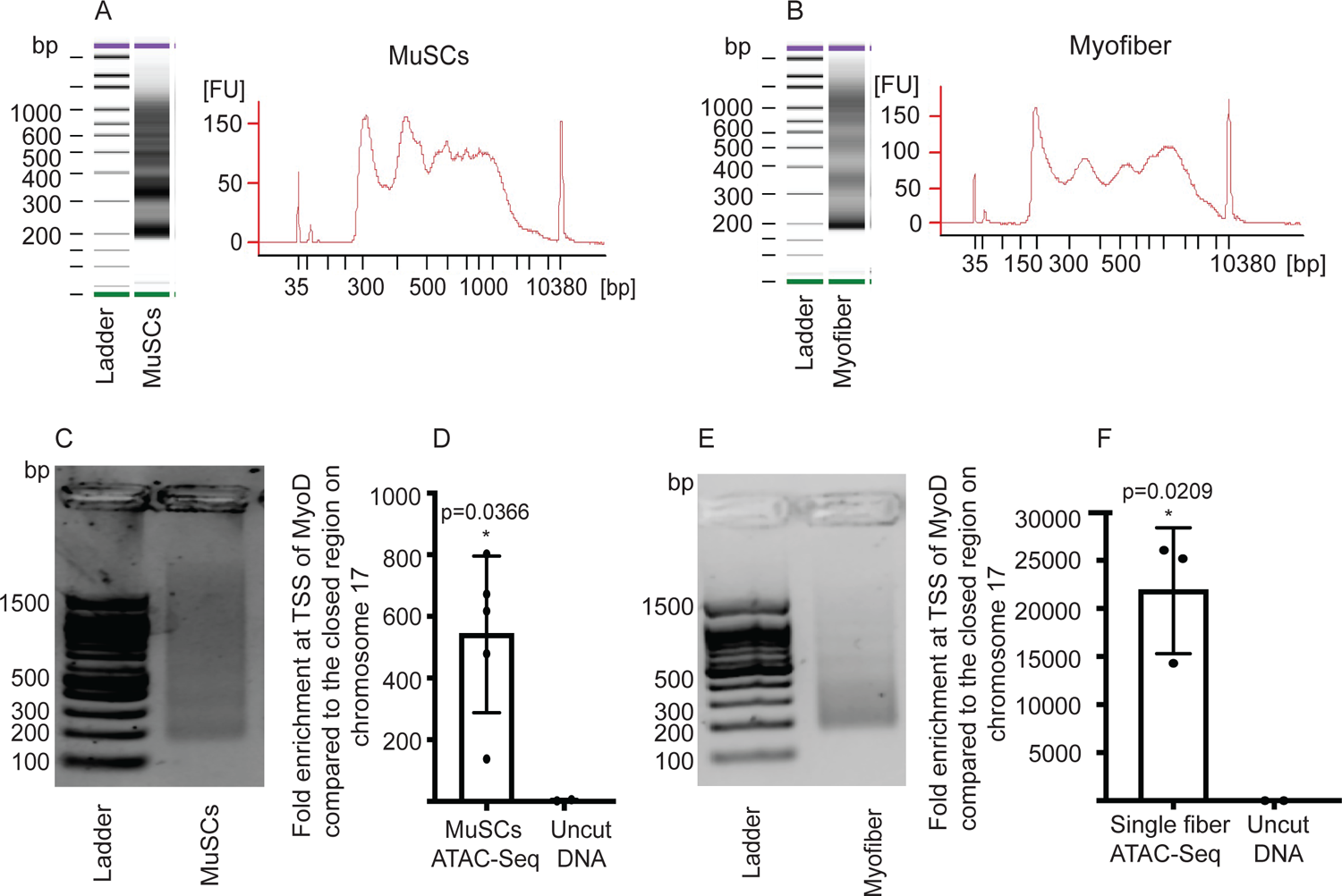
Quality control of ATAC-Seq libraries. (A) Bioanalyzer profile of an ATAC-Seq library prepared from 5000 MuSCs. (B) Example bioanalyzer profile of ATAC-Seq library prepared from a single myofiber. (C) Representative picture of a ready to sequence MuSC ATAC-seq library after size selection, visualized on an agarose gel. (D) Fold enrichment for the TSS of MyoD compared to negative control region of Chromosome 17 qE5 between MuSC ATAC-Seq libraries and untagmented DNA as seen by qPCR (n=5, 3 biological replicates and 2 technical duplicates), two-tailed t-test, error bars = ± SD. (E) Representative picture of a ready to sequence ATAC-seq on a single myofiber library after size selection, visualized on an agarose gel. (F) qPCR for the TSS of MyoD compared with a negative control region of Chromosome 17 qE5 for the ATAC-Seq libraries prepared from single myofibers (n=3, biological triplicates), two-tailed t-test, error bars = ± SD.

**Figure S2.**
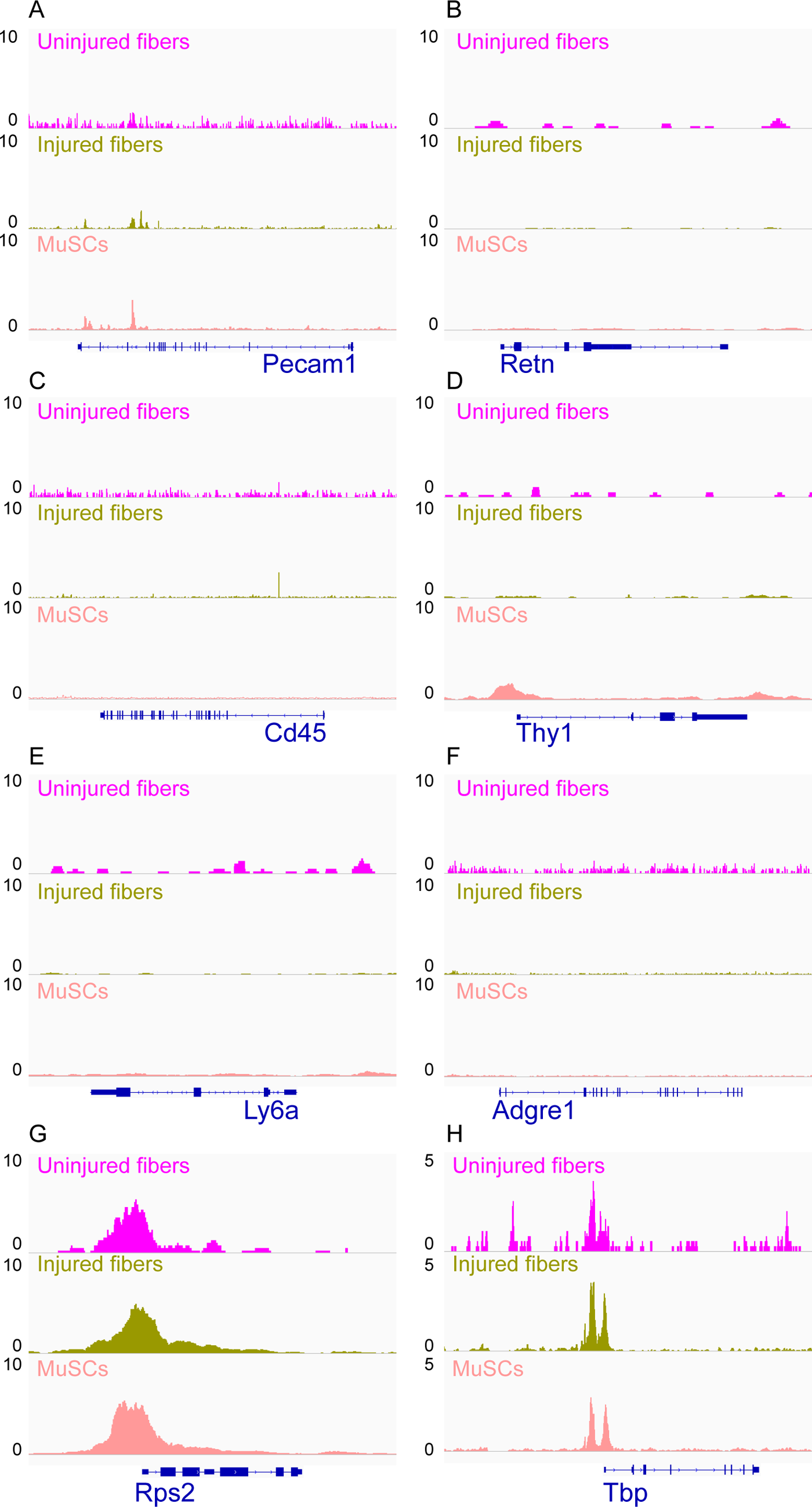
IGV snapshots of non-myogenic genes. (A) Platelet and Endothelial Cell Adhesion Molecule 1 (*Pecam1*) expressed in endothelial cells. (B) Resistin (*Retn*) as a marker of adipocytes. (C) *CD45* expressed in hematopoietic cells. (D) CD90 (*Thy1*) expressed in fibroblasts. (E) Lymphocyte antigen 6a (*Ly6a*) expressed in fibro/adipogenic progenitors (FAPs). (F) Adhesion G protein-coupled receptor E1 (*ADGRE1*) gene expressed in macrophages. (G) The housekeeping gene *RPS2* used as a positive control. (H) Housekeeping gene TATA-Box Binding protein (*Tbp*) was also used as a positive control.

**Figure S3:**
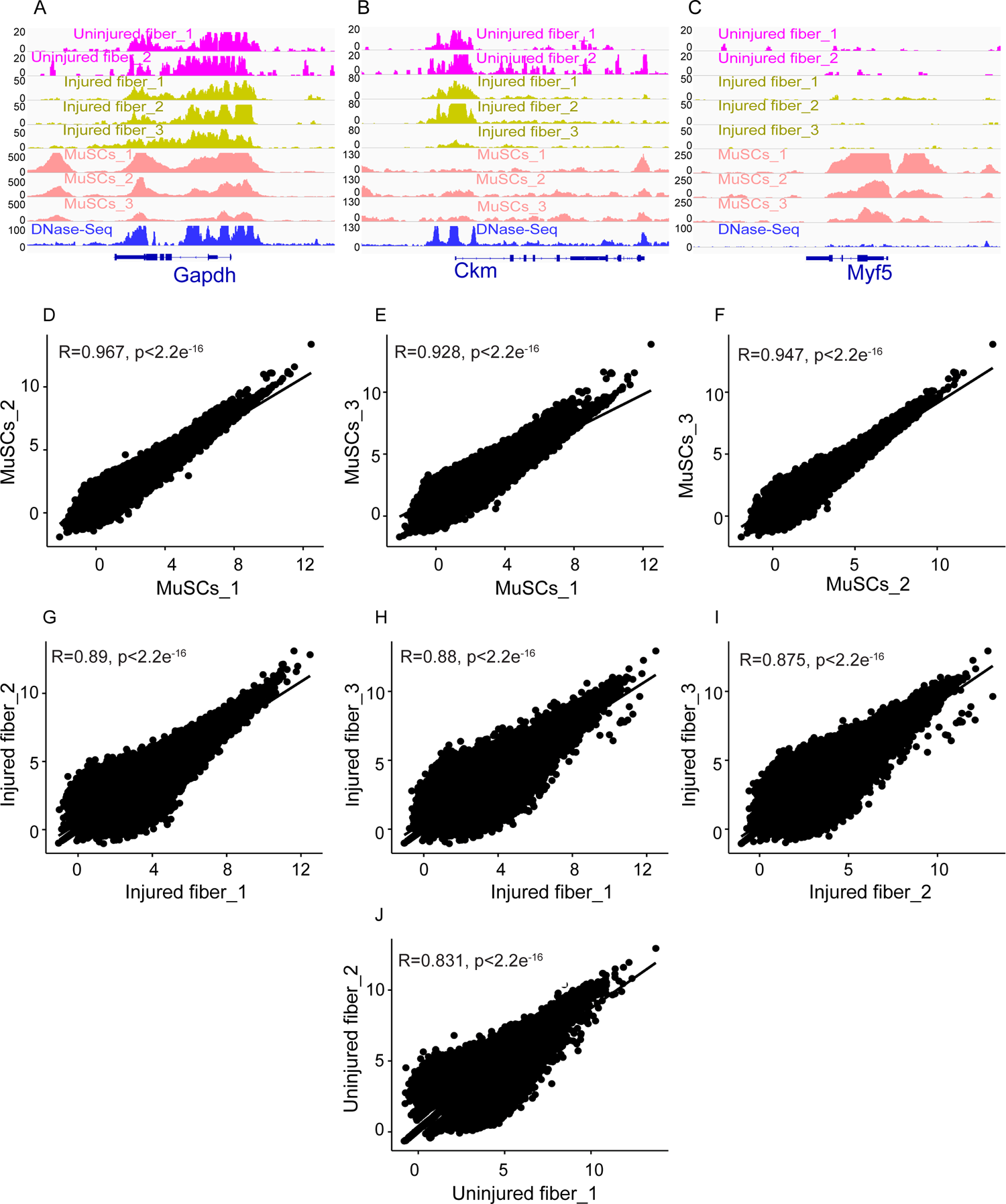
Correlation analysis between biological replicates of each condition. (A-C) IGV snapshots of genes expressed in myofibers and MuSCs for all the replicates of each condition that were pooled together for further analysis. DNase-Seq track added to demonstrate correlation of the myofiber ATAC-Seq with previously performed DNase-Seq on skeletal muscle. (A) Housekeeping gene *Gapdh*. (B) The muscle creatine kinase (*Ckm*). (C) Myogenic Factor 5 (*Myf5*). (D-J) Scatter plot showing the Pearson correlation between the replicates.

**Figure S4.**
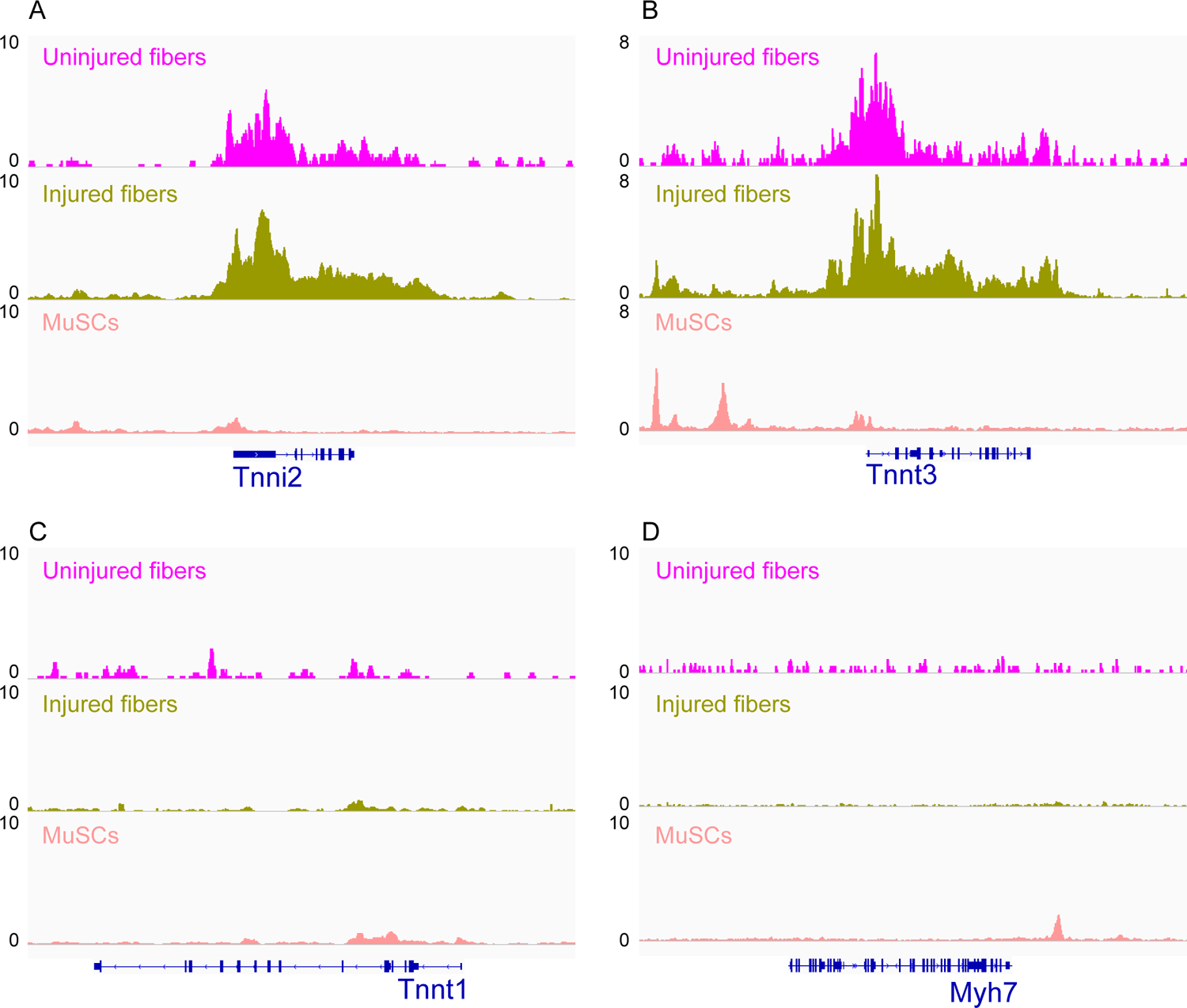
IGV snapshots of genes expressed in fast and slow muscle fiber types. (A) Troponin I2 (*Tnni2*) expressed in fast skeletal muscle fiber. (B) Troponin T3 (*Tnnt3*) expressed in fast skeletal muscle fiber. (C) Troponin T1 (*Tnnt1*) expressed in slow skeletal muscle fibers. (D) Myosin heavy chain 7 (*Myh7*) expressed in slow skeletal muscle fibers

**Figure S5:**
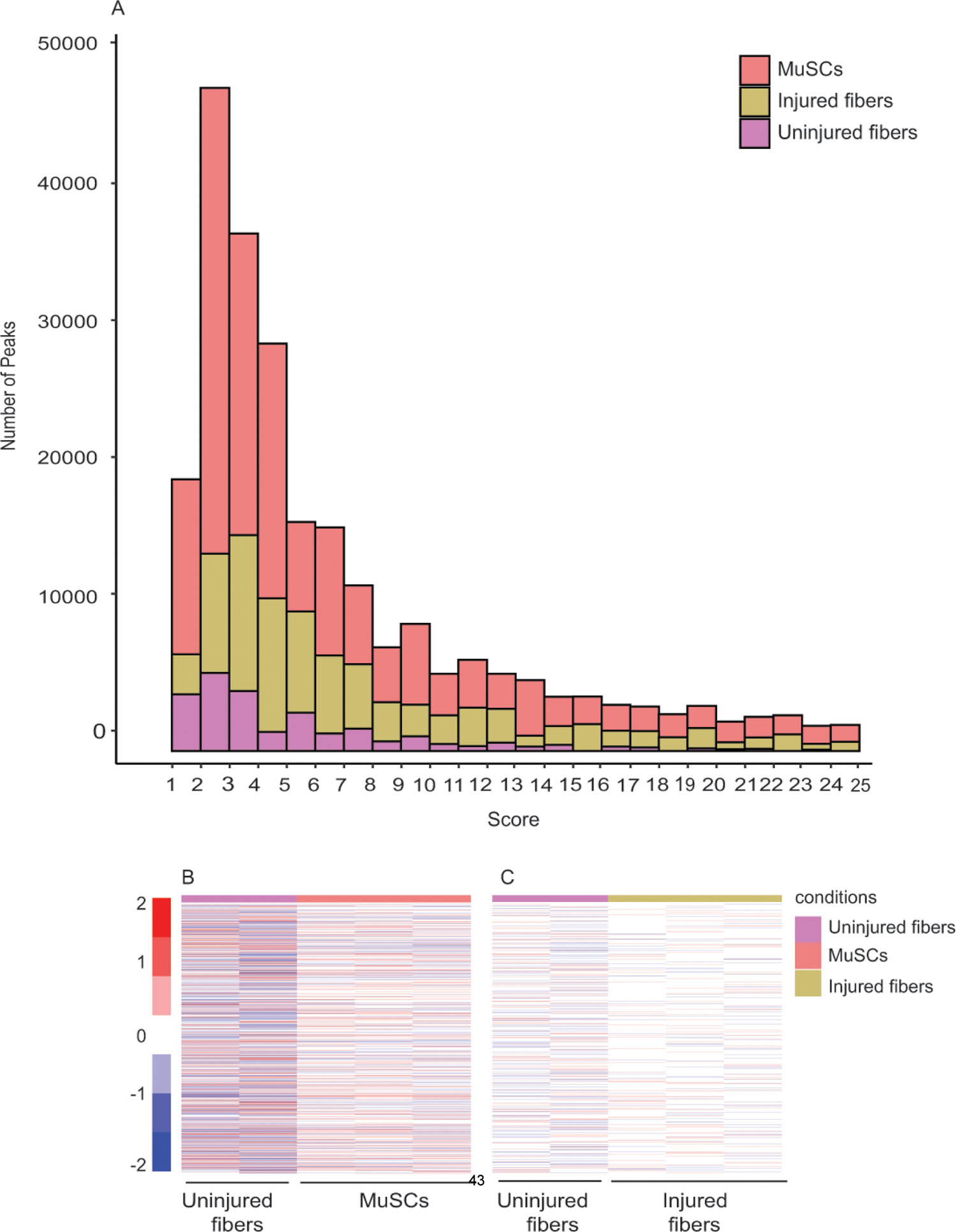
Unique peaks between different conditions indicate a distinct chromatin state for each cell type. (A) Heatmap showing the read count +/– 500 bp of the center of unique peaks to MuSCs compared to uninjured myofibers. (B) Heatmap showing the read count +/– 500 bp of the center of unique peaks to injured myofibers compared to uninjured myofibers. (C) Peak score distribution (calculated by MACS2 peak calling algorithm) for each of the different conditions. Peak score = -log10 (FDR).

**Figure S6:**
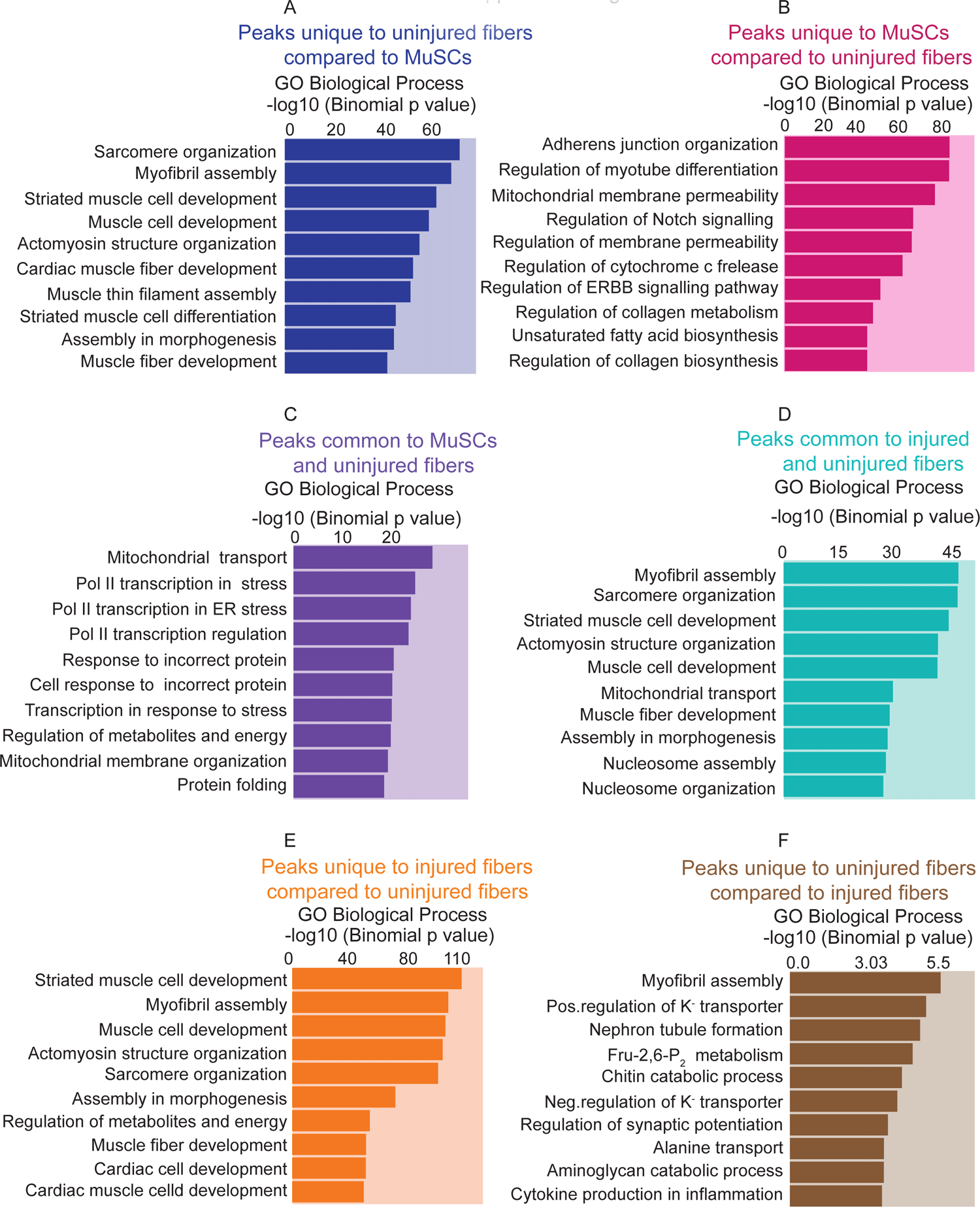
Gene Ontology analysis of unique and common peaks between conditions. (A) Gene Ontology (GO Biological Process) analysis of genes associated with unique peaks present in the uninjured myofiber compared to MuSCs, based on the proximity of the peaks to the genes. (B) GO term analysis of genes associated with unique peaks in MuSCs compared to uninjured myofibers. (C) GO term analysis of genes associated with peaks that are common between MuSCs and uninjured myofibers. (D) GO term analysis of genes associated with peaks that are common between injured and uninjured myofibers. (E) GO term analysis of genes associated with unique peaks in injured myofibers compared to uninjured myofibers. (F) GO term analysis of genes associated with unique peaks in uninjured myofibers compared to injured myofibers.

**Figure S7:**
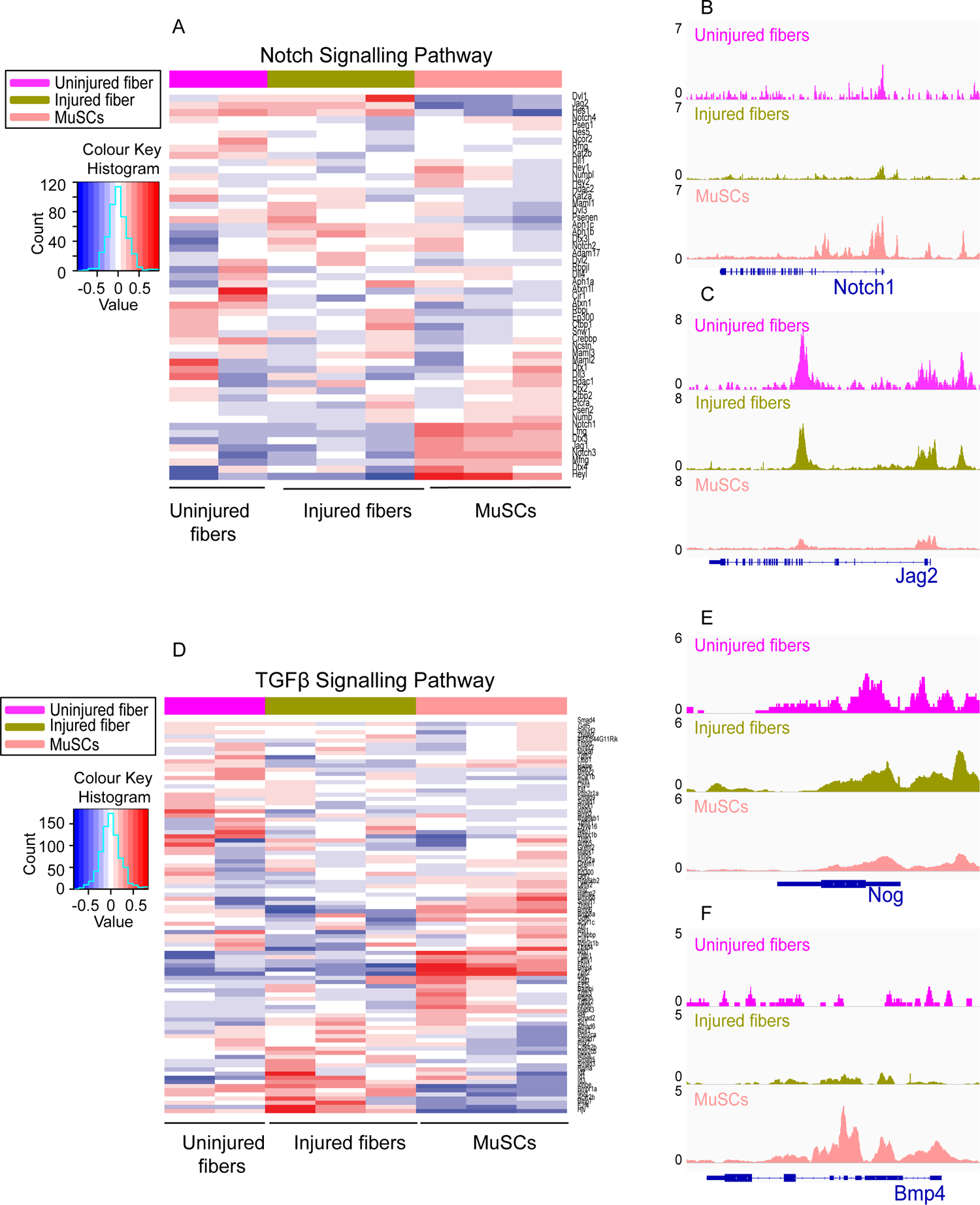
Analysis of Notch and TGFβ signalling pathways reveal differential accessibility between MuSCS and uninjured myofibers and injured and uninjured myofibers. (A)Heatmap showing genes involved in the Notch signalling pathway based on read counts of MuSCs, uninjured fibers and injured fibers, +/- 1 kb of the TSS of each gene in the pathway. (B) IGV snapshot of Notch homolog 1 (*Notch1*). (C) IGV snapshot of Protein jagged 2 (*Jag2*). (D) Heatmap showing genes involved in the TGFβ signalling pathway based on read counts of MuSCs, uninjured fibers and injured fibers, +/- 1 kb of the TSS of each gene in the pathway. (E) IGV snapshot of Noggin (*Nog*). (F) IGV snapshot of Bone morphogenetic protein (*Bmp4*).

